# Adipose tissue mitochondrial respiration in Atlantic salmon: implications for sex-dependent life-history variation

**DOI:** 10.1101/2022.07.25.501389

**Authors:** Jenni M. Prokkola, Anita Wagner, Elisabeth Koskinen, Paul V. Debes, Andrew House, Eirik R. Åsheim, Craig R. Primmer, Eija Pirinen, Tutku Aykanat

**Author notes:** Equal contribution (Jenni M. Prokkola, Anita Wagner). Corresponding author, Natural Resources Institute Finland (LUKE), Paavo Havaksen tie 3, FI-90014 Oulun yliopisto, Oulu, Finland. Author contributions: Conceptualization: JMP, AW, EP, TA Formal analysis: JMP, AW, EP, TA Funding acquisition: CRP, EP, TA Investigation: JMP, AW, EK, PVD, AH, ERÅ, CRP, EP, TA Methodology: JMP, AW, EK, PVD, CRP, EP, TA Resources: PVD, AH, ERÅ, CRP, EP, TA Supervision: JMP,EP, TA, CRP Visualisation: JMP, AW, EP, TA Writing – Original Draft Preparation: JMP, AW, EP, TA Writing – Review & Editing: JMP, AW, EK, PVD, AH, ERÅ, CRP, EP, TA.

## Abstract

Adipose tissue is essential for energy homeostasis, with mitochondria having a central role in its function. Mitochondria-mediated white adipose tissue dysfunction has been linked to several metabolic disorders in humans but surprisingly little is known about natural variation in mitochondrial function in wild animal populations, and its evolutionary significance. Early sexual maturation (low age-at-maturity) in Atlantic salmon (*Salmo salar*) is promoted by higher adiposity and has a strong genetic association with the *vgll3* locus. This makes Atlantic salmon a convenient wild model to study the potential role of mitochondria-mediated adipose tissue processes in relation to the timing of maturation. Yet, mitochondrial respiration has not been measured in the adipose tissue in fish, and the lack of data is restricting the development of informed hypotheses. Here, using 13 Atlantic salmon individuals reared in common-garden conditions, we first verified the feasibility of measuring mitochondrial respiration in the adipose tissue. As expected, the respiration level was generally low, but nonetheless we successfully quantified its biological variation in the adipose tissue. Next, we analysed the potential association of mitochondrial respiration with mitochondrial DNA (mtDNA) content, adipocyte size, sex, and the *vgll3* genotype. Despite low samples sizes, mitochondrial respiration, leak respiration, and coupling capacity (P/E ratio) were marginally significantly decreased in immature females carrying with the *vgll3* early maturation compared to the alternative genotype. Based on these results, we suggested two new hypotheses on how the coupling capacity of oxidative phosphorylation could be linked with the timing of maturation via adiposity and pave the way to study the mechanistic relationships between life-history variation and mitochondrial bioenergetics in wild populations.

## Introduction

Energy allocation dynamics are central to theories of life-history evolution. At the theoretical level and in ecological studies, white adipose tissue, which is the major energy storage organ in animals, is often viewed as an “inert” energy storage. Yet, white adipose tissue affects normal metabolic functions and reproductive fitness in animals in multiple ways (Ottaviani, Malagoli, & Franceschi, 2011). White adipose tissue (hereafter adipose tissue) is the most abundant type of adipose tissue, localising mainly to visceral and subcutaneous depots. The main function of adipose tissue is to store the dietary energy in the form of fat (triglycerides) and allocate energy to other tissues by mobilizing stored fat during fasting and high energy demand processes such as growth and sexual maturation (Norgan, 1997). The endocrine function of adipose tissue, *i*.*e*., secretion of hormones and bioactive molecules, also controls whole-body energy homeostasis and facilitates the communication between adipose tissue and other organs (Mohamed-Ali, Pinkney, & Coppack, 1998). The key functions of adipose tissue are dependent on mitochondria – the organelles that produce metabolites and energy for cells (Boudina & Graham, 2014; De Pauw, Tejerina, Raes, Keijer, & Arnould, 2009; Heinonen, Jokinen, Rissanen, & Pietilainen, 2020; Martin & Obin, 2006). Surprisingly, however, mitochondrial function in the adipose tissue is poorly studied in an evolutionary sense. For example, although the modulation of energy allocation is central to the life-history theory (e.g., van Noordwijk & de Jong, 1986), whether there is natural variation in the mitochondrial function in the adipose tissue affecting life-history trait variation and fitness is unknown.

Mitochondria are dynamic organelles – their number and efficiency to produce energy (in the form of adenosine triphosphate, ATP) are affected by the energetic status of the cell, which depends on environmental conditions and tissue type, among other factors (Kadenbach, 2003; Salin et al., 2018). Mitochondrial function can be assessed by quantifying mitochondrial respiration via oxidative phosphorylation (OXPHOS), a cascade of reactions taking place across the inner mitochondrial membrane. In OXPHOS, electrons from NADH and FADH2 are transferred through a chain of protein complexes, namely complexes I-IV, ubiquinone (Q) and cytochrome c, and finally to the electron acceptor oxygen (Fig. 1a). Simultaneously, complexes I, III, and IV pump protons into the mitochondrial intermembrane space, generating a proton gradient that drives the phosphorylation of ADP to ATP by the ATP synthase enzyme. The efficiency of mitochondrial respiration is determined by the proportion of electrons in OXPHOS that are ‘coupled’ to generating ATP (coupled respiration), *versus* proton leakage and electron slippage across the inner membrane (uncoupled respiration) (Fig 1a). Intact tissues exhibit both coupled and uncoupled OXPHOS. Highly efficient mitochondria are tightly coupled, i.e., producing more ATPs per oxygen, but uncoupled respiration is an important mechanism in adaptation to changing environmental conditions (Brand, 2005; Salin, Auer, Rey, Selman, & Metcalfe, 2015).

**Fig. 1.**
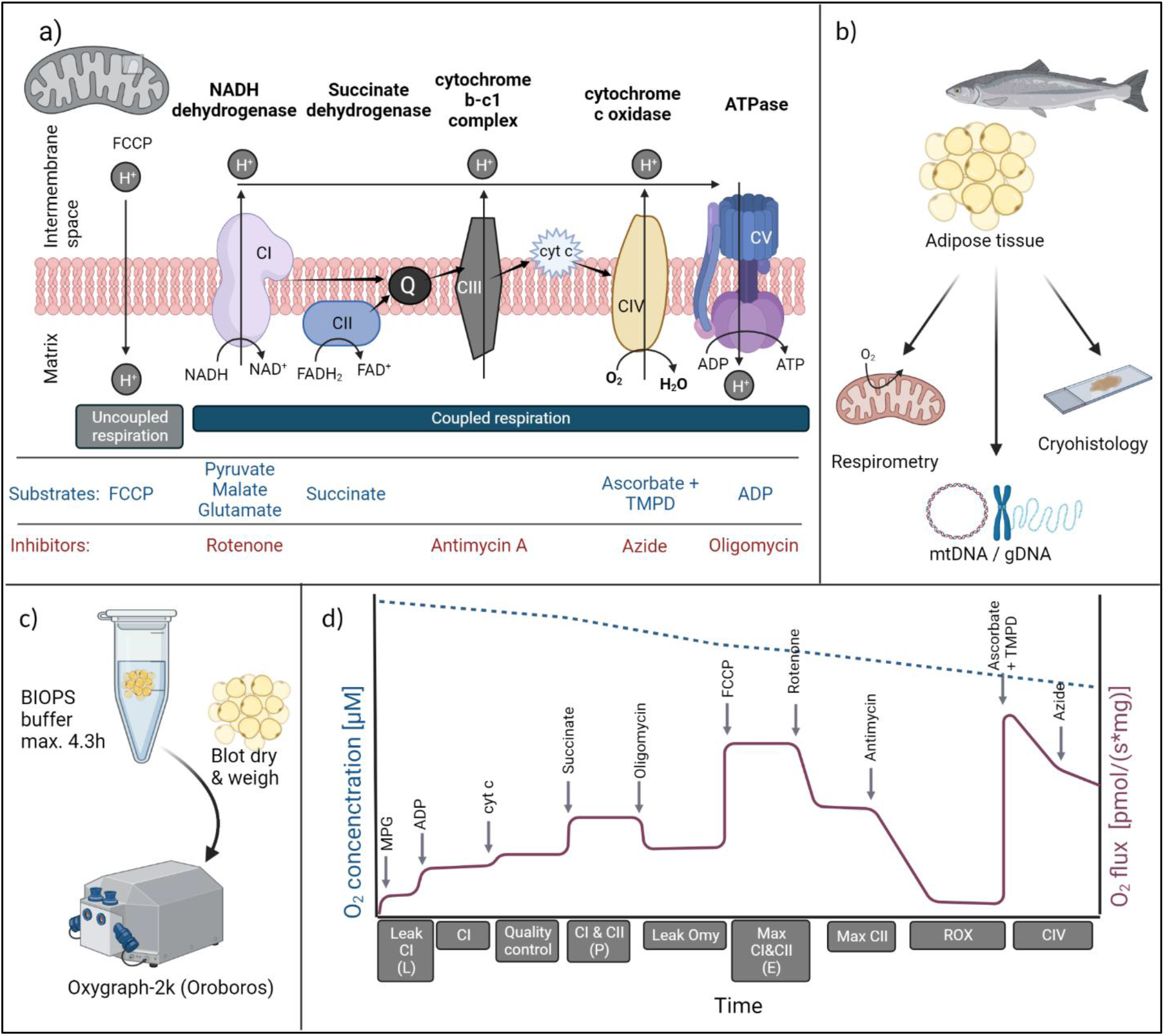
A schematic illustration of oxidative phosphorylation (OXPHOS), the study design, and the substrate-uncoupler-inhibitor respirometry protocol. (a) OXPHOS and the relevant respiration substrates and inhibitors used in this study. Electrons are shown as tapered black arrows. (b-c) The methodologies applied for visceral adipose tissue samples. (d) A schematic illustration of the injection protocol used in high-resolution respirometry with the Oxygraph 2k, with anoxygen flux curve in purple. The calculated respiration parameters are shown below the x-axis (d). MPG = malate, pyruvate, glutamate. ROX = residual oxygen flux. Figures created with BioRender.

Variation in the organisation and efficiency of mitochondria among individuals is likely under selection and adaptive (Salin, Auer, Rey, Selman, & Metcalfe, 2015; Salin et al., 2016; Hood et al., 2018; Koch et al., 2021). Within species, how variation in mitochondrial function could lead to variation in other traits is poorly understood but measuring the different stages and efficiency of mitochondrial respiration may provide a coherent framework (see e.g., Koch et al., (2021)). For example, the ability to increase ATP synthesis, which requires sufficient reserve capacity in the electron transport pathway, allows organisms to respond *via* mitochondria to changes in energetic demand and environmental stressors (Chacko et al., 2014; Sokolova, 2018). Likewise, although proton leak reduces the efficiency of mitochondria, it may restrict the production of reactive oxygen species and may thereby limit oxidative stress (Dennery, 2010). Variation in mitochondrial density and processes may thus allow individuals to respond differently to energetic demands and stressors. Despite the central role of mitochondria in the control of adipose tissue function, little is known of how adipose tissue mitochondrial activity relates to growth and body condition and affects life-history traits, such as the timing of sexual maturation. The fact that obesity in humans is associated with a significant decline in adipose tissue mitochondrial respiration (Heinonen, Jokinen, Rissanen, & Pietilainen, 2020; Jokinen, Pirnes-Karhu, Pietilainen, & Pirinen, 2017) also makes it appealing to study adipose tissue mitochondria in relation to life-history decisions in other species.

Life-history variation is largely shaped by energy allocation differences and maintained within species via evolutionary trade-offs (Lailvaux & Husak, 2014). In Atlantic salmon (*Salmo salar*), earlier maturation shortens generation time and increases the survival probability prior to reproduction compared to delayed maturation, but this comes at the expense of a smaller size at maturity, which is associated with lower fecundity (Fleming, 1998). The Atlantic salmon is an emerging wild model species to study the energetic basis of life-history adaptations for two main reasons (Mobley et al., 2021; Prokkola et al., 2022). First, faster accumulation of adipose tissue, quantified as a high condition factor is associated with earlier sexual maturation in salmon (Debes et al., 2021; House et al., 2021; Rowe, Thorpe, & Shanks, 1991). Second, a single genomic region explains a substantial amount of variation in age-at-maturity (Barson et al., 2015). The strongest candidate gene in this region, *vgll3* (Sinclair-Waters et al., 2020), is likely important for adipose tissue growth and function. In mice, for example, expression of *vgll3* was negatively correlated with adipose tissue mass and body weight (Halperin, Pan, Lusis, & Tontonoz, 2013). Likewise, in humans, variation in the *VGLL3* locus is associated with the timing of puberty (Cousminer et al., 2013; Elks et al., 2010). Finally, *vgll3* early maturation genotype is linked to a temporal increase in body condition in salmon (Debes et al., 2021). Therefore, adipose tissue energetics could provide a functional basis for variation in age-at-maturity, and subsequently, for potential evolutionary constraints (Mobley et al., 2021; Prokkola et al., 2022). Yet, the lack of information on the empirical and theoretical premises, combined with relatively challenging and costly experimental design and methodological procedures (see below, Materials and Methods), restricts statistically powerful and hypothesis-driven experiments.

In this study, we first verified the feasibility of measuring variation of mitochondrial respiration in the visceral adipose tissue – which consists almost entirely of adipocytes (Weil et al., 2012) – of Atlantic salmon. Next, we aimed to gain the first insights into preliminary associations between mitochondrial respiration and mitochondrial DNA (mtDNA) content, adipocyte size, sex, and the *vgll3* genotype, albeit using relatively small sample sizes. Based on our findings, we suggested two novel hypotheses on how oxidative phosphorylation could be linked with the timing of maturation via adiposity and pave the way for future studies of the mechanistic relationships between life-history variation and mitochondrial bioenergetics in wild populations.

## Material and Methods

### Fish rearing and sampling

The experiment was approved by the Finnish Animal Experiment Board (ESAVI/42575/2019). Samples of adipose tissue were collected during August-September 2020 from fish reared as part of another study (Åsheim et al., 2022) (here, we collected samples only from Neva population individuals from the warm temperature treatment). The fish were approximately 2 years 8 months post hatch (average mass 1 kg; Table 2). Details of fish rearing, feeding, and temperatures are shown in (Åsheim et al., 2022) until Feb 2020 (in Feb-Aug 2020, conditions largely followed those in 2019). The mean (± SD) temperature for the tanks included in this study during the sampling period was 11.7 ± 0.8 °C. Table 2 shows a summary of the number of fish used and their distribution among the experimental variables.

The fish were fasted for ∼57 h prior to sampling. One day prior to sampling, the individuals, which had been previously tagged with passive integrated transponders (PIT-tags) and identified for *vgll3* genotype and sex, were selected for sampling, anaesthetised with buffered tricaine methanosulfonate (MS-222, 0.125 g/L, sodium bicarbonate buffered), and measured for body mass (to the nearest 0.1 g) and fork length (to the nearest mm). After measurement, the fish were placed in a floating cage held inside the rearing tank until sampling the following day. The sampling was balanced in terms of sex and *vgll3* genotypes across four sampling days (from in total four tanks), on which four individuals (one female and one male of both EE and LL genotypes, referring to homozygous early and late maturation genotypes, respectively) from within the same rearing tank were captured by netting each day and euthanized with an overdose of MS-222 (0.250 g/L, sodium bicarbonate buffered). Fish were sampled from a different rearing tank between 8:40 and 11:40 AM on each day of sampling; three tanks had been reared with normal feed used in Atlantic salmon aquaculture (control), and one tank with a low-fat feed diet (details in (Åsheim et al., 2022)). Visceral adipose tissue samples were collected using sterilized equipment within ∼30 min of euthanasia and 1) stored in BIOPS (2.77 mM CaK_2_EGTA, 7.23 mM K_2_EGTA, 20 mM imidazole, 20 mM taurine, 50 mM MES hydrate, 0.5 mM DTT, 6.56 mM MgCl_2_, 5.77 mM ATP and 15 mM phosphocreatine (Canto & Garcia-Roves, 2015)), or 2) flash-frozen in liquid nitrogen and stored at -80 °C. All reagents were purchased from Sigma-Aldrich/Merck unless mentioned otherwise. Samples in BIOPS were kept on ice and transported to Meilahti campus, University of Helsinki, for mitochondrial respiration measurements (see schematic illustrations Fig. 1b-d). Due to the time-consuming nature of the measurement and requirement of fresh tissue, at most four samples were analysed each day. After sample collection, all following analyses were performed blind with respect to the diet treatment, genotype, and sex of fish.

The maturation status of the fish was determined from the appearance of gonads. All males had partially enlarged gonads indicating that the maturation process had started (the gonads of most of the remaining males in the experimental population were fully matured roughly two months after the sampling [E. Å., personal observation]), and all females had immature gonads, i.e., small eggs that covered only a small part of the body cavity, except one female that showed more advanced maturation indicated by gonad mass that was on average 30 times larger than in the other females. The maturing female was excluded from the analyses.

### High-resolution respirometry

Visceral adipose tissue stored in BIOPS buffer was used in high-resolution respiration measurements with Oxygraph-2k equipment (O2k, Oroboros Instruments, Innsbruck, Austria) (Fig.1d). Adipose tissue was dried by blotting on a tissue paper and weighed before placing it into a calibrated chamber with the Mir05 buffer (0.5 mM EGTA, 3 mM MgCl_2_, 60 mM lactobionic acid, 20 mM taurine, 10 mM KH_2_PO_4_, 20 mM HEPES, 110 mM D-sucrose, 1 g/L BSA), (Canto & Garcia-Roves, 2015) (Fig 1c). We used 28–50 mg adipose tissue in each measurement (Tables 2 and S2). The measurements were performed at 12 °C under constant stirring (750 rpm) and oxygen concentration was kept above 250 μM. Oxygen flux was calculated as means over ∼1 min within each measured parameter after a stable flux was achieved.

Substrate-uncoupler-inhibitor respirometry protocol (Fig. 1d) was used to determine LEAK (dissipative, non-OXPHOS respiration), OXPHOS (respiration coupled to phosphorylation of ADP to ATP) and ETS (electron transfer system uncoupled from the phosphorylation) respiration states. The stock solutions and final concentrations of substrates, uncoupler, and inhibitors used are presented in Table 1. To determine CI - mediated leak respiration, the NADH-pathway substrates malate, pyruvate and glutamate were added. After a stable plateau was formed, ADP at the saturating concentration was premixed with MgCl and added to induce coupled phosphorylating CI (NADH)-linked respiration. Mitochondrial outer membrane integrity was evaluated by the lack of response to the addition of cytochrome c. Next, to measure CI & CII -mediated respiration, succinate was injected to induce the electron flow through CII. After O_2_ flux was stabilised, the ATP synthase inhibitor, oligomycin, was added to determine CI&CII mediated leak respiration. To measure the maximal ETS capacity of CI&CII-linked respiration, the stepwise addition of the uncoupler carbonyl cyanide-p-trifluoromethoxyphenylhydrazone (FCCP) (Abcam) was performed. This was followed by inhibition of CI with rotenone to determine the maximal ETS capacity via CII. After blocking electron transfer with antimycin, the complex III inhibitor, residual oxygen flux (ROX) was quantified. Complex IV (CIV) activity was determined by adding N,N,N’,N’-Tetramethyl-p-phenylenediamine dihydrochloride (TMPD) as a substrate in conjunction with the reducing agent ascorbate. Complex IV was inhibited with sodium azide to correct previous results for the chemical auto-oxidation of reagents (Djafarzadeh & Jakob, 2017). Oxygen flux was quantified using the DatLab analysis software (Canto & Garcia-Roves, 2015). ROX was subtracted from all except CIV-related values. The delay until the respiration measurements started after the fish were euthanized ranged from 1.8 to 4.3 h but this was not correlated with oxygen flux during the measurements (Fig. S1).

**Table 1.**
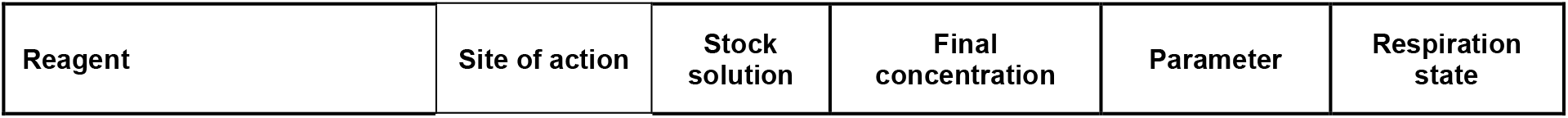
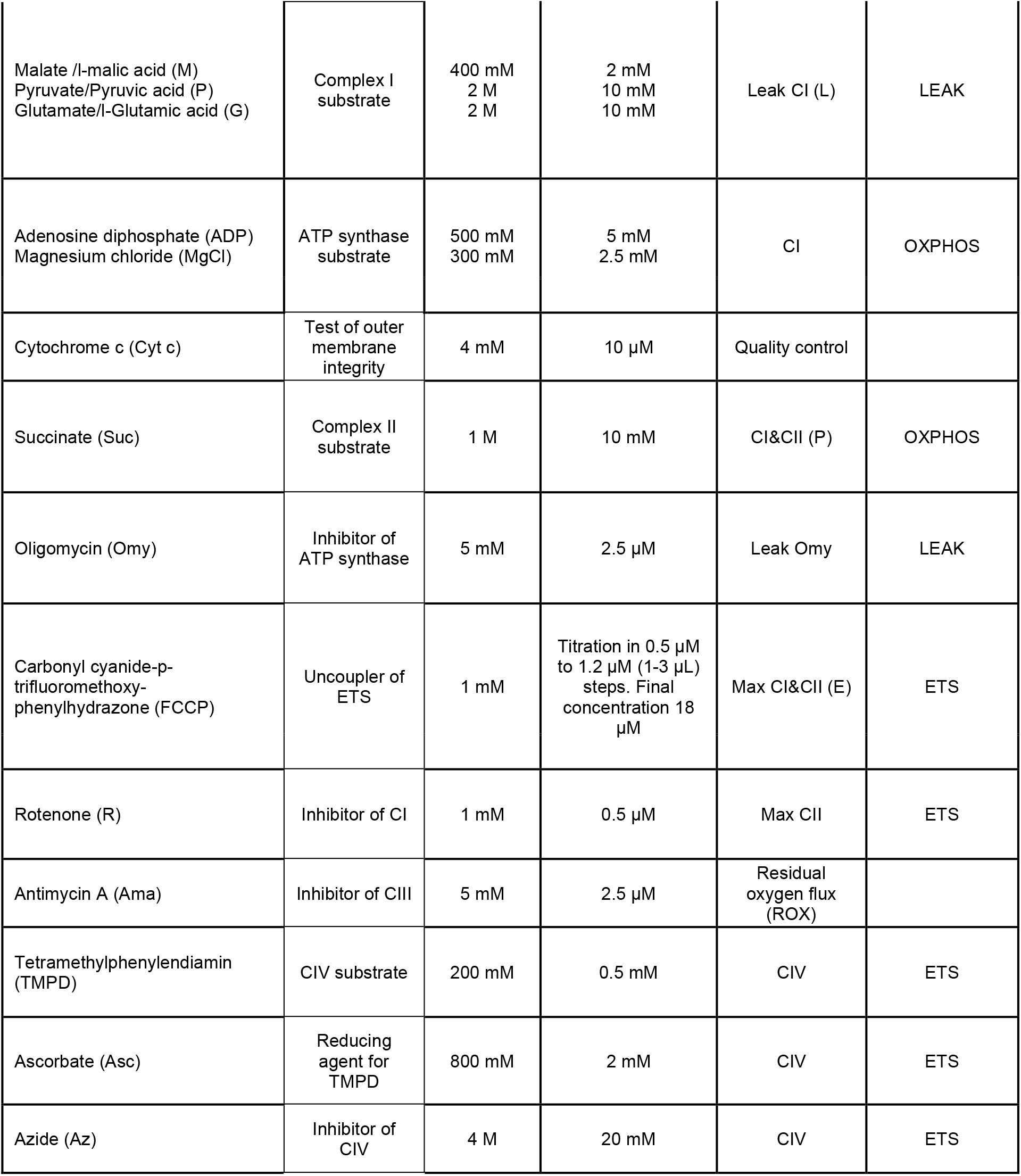
The used reagent concentrations and their respective respiration parameters and states.

We calculated mitochondrial respiration coefficients from tissue mass -normalised parameters shown in Table 1 and Fig. 1d as follows: *NADH pathway cofactor (NADH)*: CI / CI&CII, indicating the proportion of CI respiration from CI & CII respiration; *Succinate pathway cofactor (Succ)*: 1 - CI / CI&CII, indicating the proportion of CII respiration from CI & CII respiration; *Coupling efficiency (CoupEff)*: (CI&CII - Leak Omy) / CI&CII, indicating the proportion of ATP synthesis -linked respiration from CI & CII respiration; *Coupling control ratio (L/P):* Leak CI / CI&CII, indicating the proportion of CI-linked proton leak from CI & CII respiration; and *Coupling capacity (P/E)*: CI&CII / max CI&CII, indicating the proportion of phosphorylating, coupled CI & CII mediated respiration from non-phosphorylating, uncoupled CI & CII mediated respiration.

### Measurements of mitochondrial DNA content

We quantified mtDNA amount relative to nuclear genomic DNA (gDNA) as a proxy for mitochondrial content per cell (Robin & Wong, 1988) to understand whether mitochondrial respiration variation was explained by differences in mitochondrial number. The tissue used in this procedure was collected at the same time as the sample used in the high-resolution respiration measurements. First, DNA was extracted using a slightly modified conventional phenol-chloroform extraction. Phenol chloroform extraction conserves different sizes of DNA relatively similarly compared to silica column-based extraction protocols, hence exhibiting higher reproducibility when quantifying mtDNA to gDNA ratio (Guo, Jiang, Bhasin, Khan, & Swerdlow, 2009). Briefly, approximately 100 mg of adipose tissue and a 7 mm metal bead were placed in a 2 mL centrifuge tube containing 0.5 mL PBS buffer (pH 7.4) and homogenised in TissueLyser II (Qiagen) for 2 minutes at 25 Hz. The homogenate was then incubated overnight at 37 °C in 0.6 mL lysis buffer (10 mM Tris–HCl, 1 mM EDTA, and 0.1% SDS, pH 8.0) with 7 µL proteinase K solution (20 mg/mL), and then treated with RNAase (0.2 mg/mL) for 30 min at 37 °C. Next the solution was treated with 600 μL phenol (VWR, Tris-buffered, pH=6.6) twice, followed by two 500 μL chloroform isoamyl alcohol (24:1) washes. A 1:10 volume of sodium acetate (3 M, pH 5.2) and 2.5 volumes of ice-cold ethanol were then added to the aqueous phase and samples were incubated for 30 min at -20 °C. The precipitated DNA was then washed twice with 70 % ethanol, air dried for 30 min, and resuspended in 50 μL TE solution (5 mM Tris, 0.1 mM EDTA, pH 8.0) for 15 min at 37 °C. DNA quality and quantity were assessed using Nanodrop 2000 (Thermo Scientific) and DNA concentration was adjusted to 5 ng/μL. All samples in the analyses had 260/280 ratios greater than 1.80 (average = 1.90, SD = 0.05), indicating high purity. DNA was extracted in two batches for n = 12, and n = 8 samples, respectively, where four samples were extracted twice to control for batch effects (totalling 16 samples); these samples were run in six replicate reactions, while the others were run in triplicates.

Next, quantitative real-time PCR (qPCR) was performed using a double-stranded DNA-binding dye as a reporter (HOT FIREPol EvaGreen qPCR Supermix, Solis Biodyne), in a 384-plate format using a Biorad CFX384 C1000 thermal cycler. We targeted two mitochondrial (*16s* and *cytb)*, and two nuclear (*app, EF1a*) genomic regions using a double-stranded DNA-binding dye (eva green) as a reporter. Primers specific to these genomic regions were designed with Primer-BLAST (Ye et al., 2012), where the melting temperature (Tm) and product size were adjusted to 59–61 °C, and 90– 150 bp, respectively (Table A1).

The qPCRs were run for 45 cycles at 95 °C, 58 °C, and 72 °C for 15, 20, and 20 s, respectively, following initial denaturation at 95 °C for 12 min. The volume of reactions was 10 μL, with 2 μL 5x HOT FIREPol EvaGreen qPCR Supermix (Solis Biodyne, Tartu, Estonia), 150 nM of each primer, and 5 μL diluted DNA (total 1.67 ng). Samples were analysed in triplicates for each marker.

The relative mtDNA to gDNA amount, where gDNA amount was first divided by two to account for two copies of gDNA per cell, was then quantified from mean expression across replicate reactions using an efficiency-corrected method (Pfaffl, 2001). Ct values and PCR efficiencies were calculated using LinregPCR (Ruijter et al., 2009), where baseline correction and window-of-linearity analysis (the cycles with the highest linear correlation) was performed separately for each well and the efficiency of each primer set was calculated as the mean of individual wells’ efficiency.

### Measurements of adipocyte size

Flash-frozen samples of visceral adipose tissue were sectioned to assess whether adipocyte size correlated with mitochondrial respiration. Samples were embedded in the Optimal Cutting Temperature compound and sectioned to 20–30 µm at 50 °C using a cryostat (Leica CM 3050S) and stained with haematoxylin & eosin using Sakuras Tissue-Tek DRS 2000 at the Department of Anatomy, University of Helsinki following a standard protocol. In brief, cryosections were fixed with ice-cold acetone for 10 min on ice, air dried for 30 min, washed twice with PBS for 1 min, incubated in PBS for 10 min, washed in water for 1 min, stained with Mayer’s hemalum solution (Merck) for 5 min, washed with tap water for 1 min, incubated in distilled water for 1 min, stained with May-grunwald’s eosin-methylene blue solution modified (Merck) for 3 min, and washed in tap water for 1 min, and twice each in 96 % ethanol for 20 s, Abs ethanol 5 min, and Xylene 2 min. Finally, slides were covered with Pertex mounting medium (Histolab).

The slides were imaged using 20x magnification with extended plane, using 3DHISTECH Pannoramic 250 FLASH II digital slide scanner. Images were inspected in Qupath v. 0.3.0 (Bankhead et al., 2017) and 1–2 regions containing visceral adipose tissue from each image were saved in tiff format. Tiff-images were analysed in ImageJ to measure the size of particles with a size of 1000–100000 and circularity of 0.2–1.00 as size and shape filters (used macro in Supplemental Material). The selected cells were visually inspected to exclude selections that were merged of multiple cells, were stretched (during sectioning) or had a very irregular shape because of excess dye. The remaining selected cells were measured (area in µm^2^) (see Fig. 2a for a representative image). The number of cells identified with this method ranged from 40 to 306 between individuals due to the variable size and quality of the sections. However, the number of measured cells was not correlated with mean (Spearman-rho = -0.24, p = 0.426) or median (Spearman-rho = -0.26, p = 0.388) adipocyte size, indicating that there was no size bias due to the number of cells measured (Fig. S2).

**Fig. 2.**
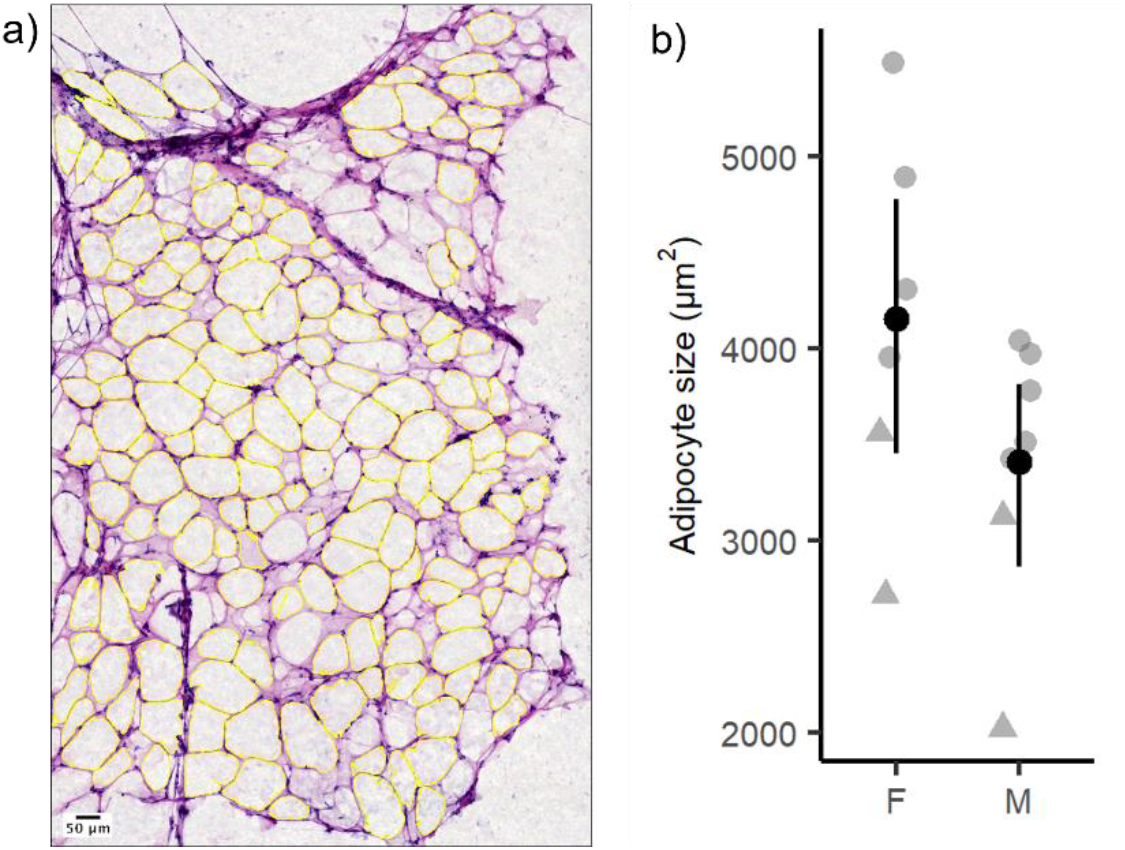
Atlantic salmon adipocyte morphology. (a) Light microscope image of a representative H&E-stained cryosection of visceral adipose tissue. Measured cells highlighted in yellow (see Material & Methods section Measurement of adipocyte size). Fifty µm scale bars shown on the image, magnification 20X. (b) Mean adipocyte size (black points) ± 95 % bootstrapped confidence intervals in Atlantic salmon females and males (n_fem_ = 6, n_male_ = 7) (Wilcoxon rank sum test, Table 2, p = 0.175). Grey symbols show individual data points: circles = control, triangles = low fat.

### Data analysis

The size, *vgll3* genotype and respiration data for each individual are provided in Table S2. Respiration was successfully measured from adipose tissues of 14 individuals (nine females, five males) after data from two males were omitted due to abnormal noise in the data (both with the early maturation *vgll3* genotype). Leak CI was not determined for three individuals due to abnormal fluxes. Cryosections for adipocyte size measurements were obtained from 14 individuals (seven of each sex, after two females were omitted from the analysis due to low section quality). Two males without respiration data were included in the adipocyte size measurements. One female that displayed stronger maturation than all others was omitted from the dataset, since salmon consume the energy stored in adipose tissue during maturation, hence the maturation status likely affects the results (Rowe et al., 1991). Given that all the remaining females in this study were immature, and all males were maturing, sex effects are confounded with maturation status throughout the study, which is taken into account when interpreting the result.

Oxygen fluxes were normalised to tissue mass as well as to both tissue mass and mtDNA amount. To compare the respiration data of individuals with different *vgll3* genotypes, we focussed on data from females, as we only obtained data from two early maturation (*vgll3*\*EE) -genotype males.

The data were analysed in the R software environment (R Core Team, 2019), and visualised using ggplot2 (Wickham, 2009). We tested the statistical significance of feed treatment and sex effects and of genotype effects in females using non-parametric Wilcoxon rank sum tests and Spearman’s rank correlations to avoid violations of linear model assumptions due to low sample sizes. In line with the exploratory nature of this study, multiple-test correction, which is also restrictively conservative at small samples sizes, was not employed.

## Results

### Data overview and feed treatment effects

Because samples were obtained from salmon under different feeding treatments, we first determined the effect of diet fat content on fish phenotypes. No effects of feeding treatment were detected on fish morphology (Table 2) or respiration traits (see Fig. 2b and Fig. 3-4, where feed treatments are shown with different symbols, and Table S3) except adipocytes were significantly larger in salmon reared under a control diet than a low-fat diet, as expected (Table 2). Data from the two feeding treatments were combined to improve statistical power of the subsequent analyses.

**Table 2.**
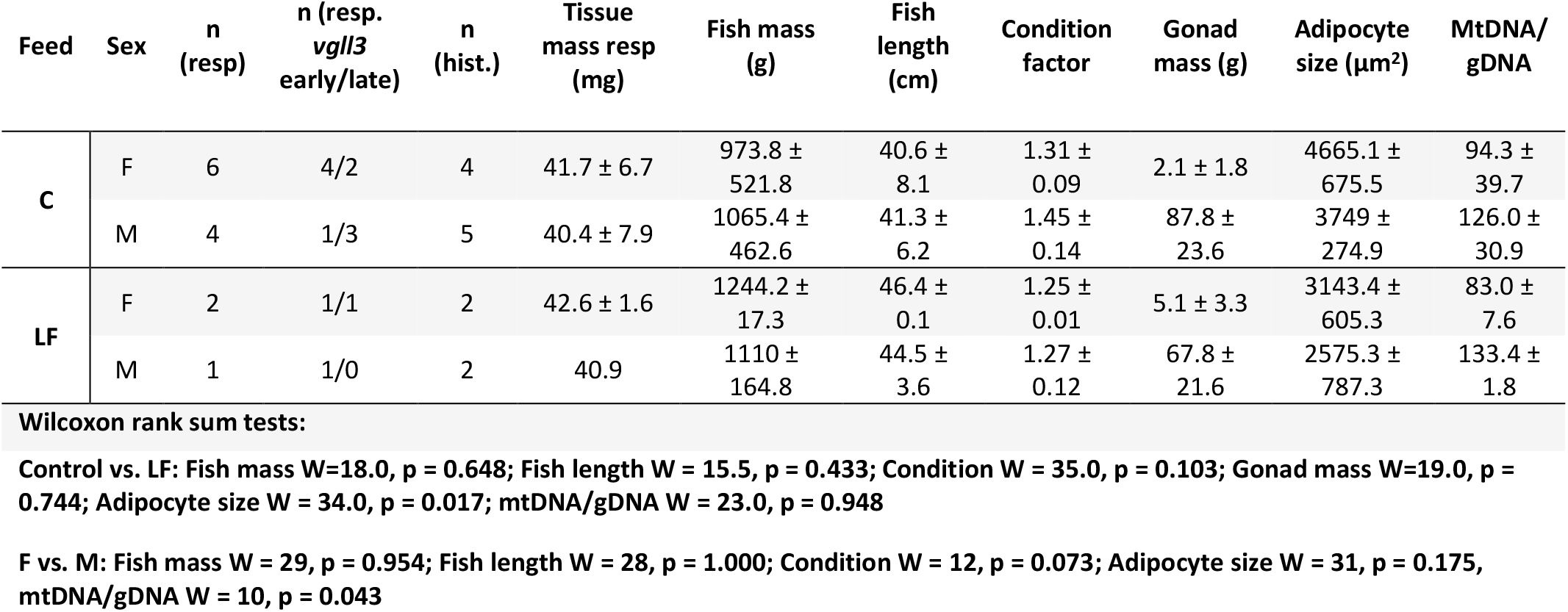
Summary of final data sets and fish phenotypes (mean ± SD) from control (C) and low fat (LF) feed treatments. All differences between treatments were non-significant, apart from adipocyte size (results shown in footnotes). Resp = mitochondrial respiration, hist = cryohistology. Full data is provided in Table S1.

| Feed | Sex | n<br>(resp) | n (resp.<br>vgll3<br>early/late) | n<br>(hist.) | Tissue<br>mass resp<br>(mg) | Fish mass<br>(g) | Fish<br>length<br>(cm) | Condition<br>factor | Gonad<br>mass (g) | Adipocyte<br>size ($\mu\text{m}^2$) | MtDNA/<br>gDNA |
| --- | --- | --- | --- | --- | --- | --- | --- | --- | --- | --- | --- |
| C | F | 6 | 4/2 | 4 | 41.7 ± 6.7 | 973.8 ±<br>521.8 | 40.6 ±<br>8.1 | 1.31 ±<br>0.09 | 2.1 ± 1.8 | 4665.1 ±<br>675.5 | 94.3 ±<br>39.7 |
| | M | 4 | 1/3 | 5 | 40.4 ± 7.9 | 1065.4 ±<br>462.6 | 41.3 ±<br>6.2 | 1.45 ±<br>0.14 | 87.8 ±<br>23.6 | 3749 ±<br>274.9 | 126.0 ±<br>30.9 |
| LF | F | 2 | 1/1 | 2 | 42.6 ± 1.6 | 1244.2 ±<br>17.3 | 46.4 ±<br>0.1 | 1.25 ±<br>0.01 | 5.1 ± 3.3 | 3143.4 ±<br>605.3 | 83.0 ±<br>7.6 |
| | M | 1 | 1/0 | 2 | 40.9 | 1110 ±<br>164.8 | 44.5 ±<br>3.6 | 1.27 ±<br>0.12 | 67.8 ±<br>21.6 | 2575.3 ±<br>787.3 | 133.4 ±<br>1.8 |

**Fig. 3.**
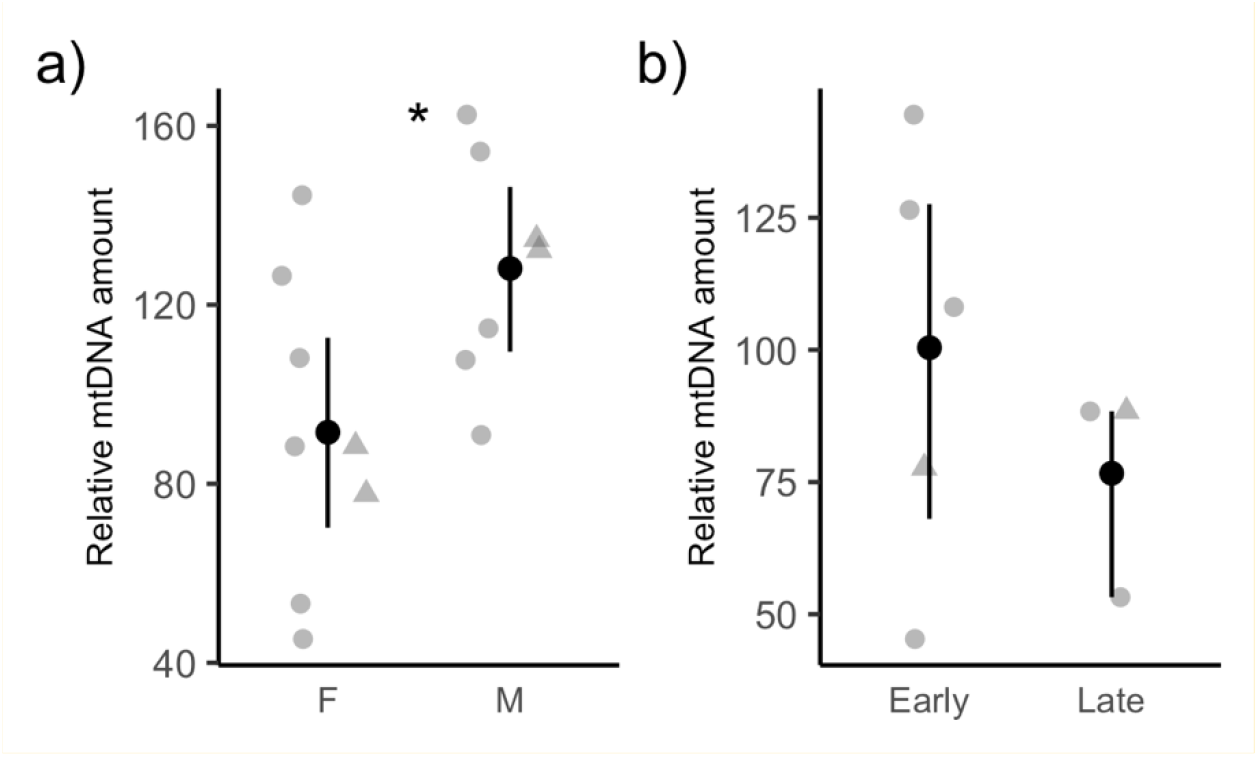
Relative mtDNA amount means ± 95 % bootstrapped confidence intervals in (a) different sexes and (b) only females carrying *vgll3* genotypes related to either early or late maturation. n_fem_ = 8, n_male_ = 7, n_early_ = 5, n_late_ = 3. Wilcoxon rank sum tests: (a) W = 10, p = 0.043 (asterisk), (b) W = 10, p = 0.551. Grey points show individual data points: circles = control, triangles = low fat.

### Sex differences

All males had enlarged gonads verifying that maturation had been initiated. Females were immature (see also Materials and methods). Males had a higher relative mtDNA amount than females (Table 2; Fig. 3a), but mitochondrial respiration did not differ between the sexes after the data were normalised either with tissue mass, or tissue mass and relative mtDNA amount (Table 3; Fig. 4a-b). This result was corroborated also by a lack of difference between the sexes in mitochondrial respiratory coefficients (Fig. S3). Interestingly, in both sexes CI-mediated respiration (NADH pathway cofactor) was substantially higher than CII-mediated respiration (succinate pathway cofactor) (Fig. S3), suggesting a generally higher preference for NADH-driven electron transfer in salmon visceral adipose tissue. There were no sex differences in any other morphological phenotypes measured nor in adipocyte size (Table 2; Fig. 2b).

**Table 3.** Results of Wilcoxon rank sum tests of sex- and genotype effects on mitochondrial respiration parameters normalised with tissue mass, or with tissue mass and mtDNA amount. For n see Tables 2 and 4. Significant p-values in bold.

| Comparison | Variable | Mass corrected |  | Mass & mtDNA corrected |  |
| --- | --- | --- | --- | --- | --- |
|  |  | W | p | W | p |
| Female vs. Male | Leak CI | 12 | 1.000 | 17 | 0.352 |
|  | CI | 12 | 0.272 | 27 | 0.354 |
|  | CI&CII | 14 | 0.421 | 27 | 0.354 |
|  | Leak Omy | 15 | 0.510 | 27 | 0.354 |
|  | Max CI&CII | 16 | 0.608 | 28 | 0.284 |
|  | Max CII | 11 | 0.213 | 29 | 0.222 |
|  | CIV | 27 | 0.341 | 33 | 0.065 |
| *Vgll3* Early vs. Late | Leak CI | 4 | 1.000 | 3 | 0.700 |
|  | CI | **0** | **0.036** | **0** | **0.036** |
|  | CI&CII | **0** | **0.036** | 2 | 0.143 |
|  | Leak Omy | **0** | **0.036** | 3 | 0.250 |
|  | Max CI&CII | 4 | 0.371 | 2 | 0.143 |
|  | Max CII | 4 | 0.371 | 2 | 0.143 |
|  | CIV | 8 | 1.000 | 6 | 0.786 |

**Fig. 4.**
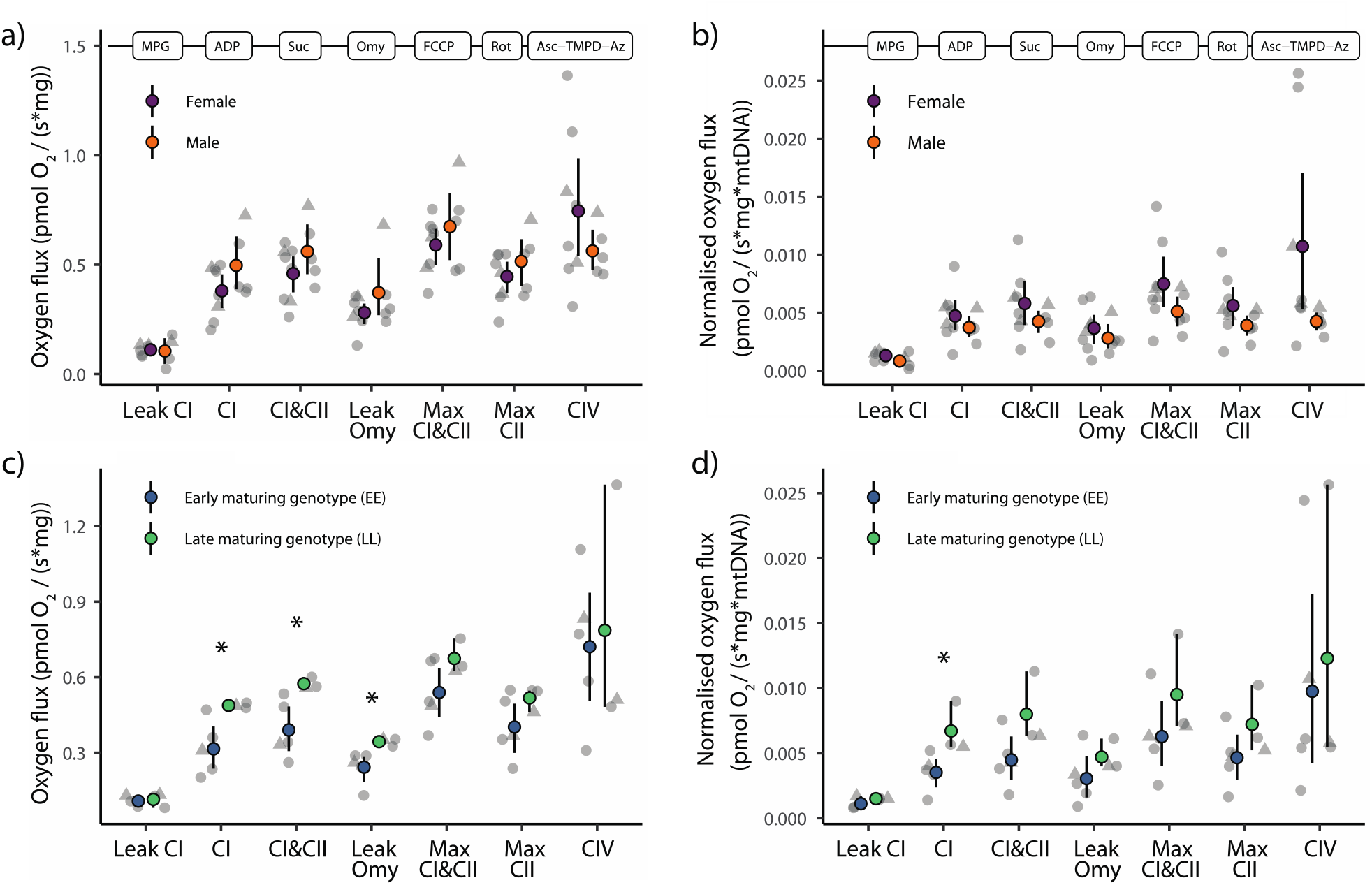
Mitochondrial respiration in Atlantic salmon adipose tissue across sexes (a,b) and *vgll3* genotypes (c,d). (a) and (c) mitochondrial respiration normalised to tissue mass, (b) and (d) mitochondrial respiration normalised to mtDNA amount in both sexes and genotypes. n_fem_ = 8, n_male_ = 5, n_early_ = 5, n_late_ = 3. Added compounds for respiration measurements are shown in chronological order over the plots (see Fig. 1d). Coloured points show means with bootstrapped 95% confidence intervals. Asterisks in (c) and (d): p = 0.036 (Wilcoxon rank sum test, Table 3). Grey points show individual data points: circles = control, triangles = low fat.

### Vgll3 effects on mitochondrial traits

To explore the relationship between *vgll3* genotypes and mitochondrial respiration, we focussed only on female salmon (Table 4) due to the limited availability of data from males with *vgll3*\*EE genotype. We found that CI- and CI&CII -mediated respiration were significantly higher in individuals with the late maturation genotype compared to those with the early maturation genotype of *vgll3* (p = 0.036; Table 3; Fig. 4c). In line with the higher respiration, CI&CII -linked respiratory leak (Leak Omy) was significantly elevated in individuals with the late maturation genotype (p = 0.036; Table 3; Fig. 4c). Consequently, there was no difference between the genotypes in ATP synthesis-linked respiration rate, i.e., Leak Omy subtracted from CI&CII (mean (bootstrapped 95% C.I.): Early = 0.148 (0.0761, 0.218) and Late = 0.230 (0.207, 0.275), Wilcoxon rank sum test W = 4, p = 0.371). We then asked whether the genotype effects were mediated by the mtDNA amount – as a proxy of mitochondrial density – and found no significant differences (p = 0.551; Fig. 3b). Finally, when respiration was normalised to mtDNA amount within the same individuals, the differences were insignificant in CI&CII-mediated respiration but remained significant in CI-mediated respiration (p = 0.036; Table 3; Fig. 4d). There were no significant effects of genotype on the other respiration traits (Table 3; Fig. 4c, d). *Vgll3* genotype effects were also mostly absent in respiration coefficients. However, the coupling capacity, P/E, of individuals carrying the late maturation genotype was marginally higher than of those carrying the early maturation genotype (p = 0.071, Fig. 5).

**Table 4.** Summary of female salmon phenotypes (mean ± SD) across early maturation (vgll3*EE) and late maturation (vgll3*LL) genotypes. No significant differences between genotypes (see footnotes). Resp = mitochondrial respiration.

| Genotype | n (resp) | Fish mass (g) | Fish length (cm) | Condition factor | Gonad mass (g) | Adipocyte size (μm) | MtDNA/gDNA |
| --- | --- | --- | --- | --- | --- | --- | --- |
| Early | 5 | 1135.4 ± 481 | 43.2 ± 7.4 | 1.31 ± 0.10 | 3.7 ± 2.7 | 4354.1 ± 1195.2 | 100.4 ± 39.5 |
| Late | 3 | 884.8 ± 462.8 | 40.2 ± 8.3 | 1.26 ± 0.01 | 1.6 ± 1.1 | 3765.5 ± 274.4 | 76.7 ± 20.3 |

Wilcoxon rank sum tests
Early vs. Late: Fish mass $W = 10$ , $p = 0.551$ ; Fish length $W = 10$ , $p = 0.551$ ; Condition $W = 9$ , $p = 0.766$ ; Gonad mass $W = 12$ , $p = 0.233$ ; Adipocyte size $W = 6$ , $p = 0.487$ , mtDNA/gDNA $W = 10$ , $p = 0.551$ .

**Fig 5.**
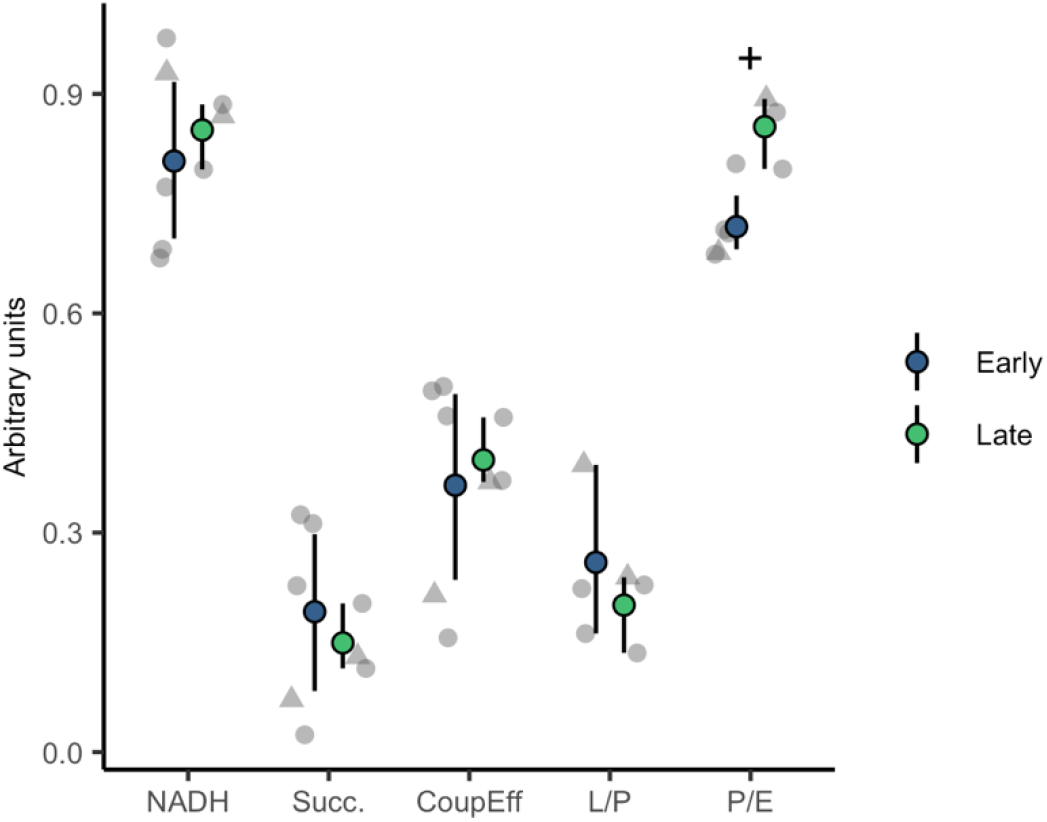
Respiration coefficients in female salmon with different vgll3 genotypes. The plus sign shows marginally higher coupling capacity (P/E) in the late than early maturation genotype (W = 1, p = 0.071, Wilcoxon rank sum test, n_early_ = 5, n_late_ = 3). Coloured points show means with bootstrapped 95% confidence intervals. Grey symbols show individual data points: circles = control, triangles = low fat treatment.

### Correlations among fish phenotypes and mitochondrial respiration

We calculated Spearman’s rank correlations between mitochondrial respiration values and fish phenotypic data, including body mass, condition factor, relative mtDNA amount and adipocyte size. There was a marginally significant negative correlation between mtDNA amount and CIV -mediated respiration (rho = -0.49, p = 0.089, Fig. S4a), but none of the other correlations were significant (p > 0.1) (Table S4).

## Discussion

Mapping the molecular and physiological basis of life-history variation is one of the key goals of evolutionary biology. Life-history diversity within and among species is ultimately determined by energy use (Lailvaux & Husak, 2014), in which adipose tissue and mitochondria functions have a central role. In this study, we provide a workflow that integrates adipose tissue mitochondrial respiration into the life-history theory framework in Atlantic salmon. Our results suggest that mitochondrial phenotypes exhibit variation associated with different backgrounds (i.e., sex/maturation, and age-at-maturity genotype). Consequently, we propose that energy generation and dissipation in the adipose tissue may have a role in the physiological determination of age at maturity via the *vgll3* genomic region. As some of the associations we observed were significant, we use the results to build novel, informed hypotheses on the potential mechanistic links between *vgll3* genotype and adipose tissue growth. Our results guide future work to test these hypotheses with larger experimental designs and point to the potential value of integrating adipose tissue mitochondrial phenotypes with life-history variation in the wild.

Overall, we verified the feasibility of measuring visceral adipose tissue mitochondrial respiration in large-sized (1 kg) Atlantic salmon. The respiration rates increased with the addition of substrates for CI and CII -mediated electron transfer, as expected. We found that visceral adipose tissue mitochondria relied more on NADH-driven CI-mediated than succinate-driven CII-mediated respiration (Fig. 4 & 5). Since NADH is generated via many different cellular processes including glycolysis, citric acid cycle and fatty acid oxidation, the activity of these cellular processes can regulate adipose tissue OXPHOS. The respiration rates were much lower than what had been earlier reported in salmonids from aerobically more active tissues, such as muscle or intestine (Brijs et al., 2017; K. Salin et al., 2016), but this was an expected outcome since adipose tissue is mostly composed of lipids (Nanton et al., 2007). Despite the lower rate of mitochondrial respiration in adipose tissue, the L/P ratio indicated that approximately 30% of mitochondrial respiration was related to proton leak, which is similar to the levels reported previously in fish intestine (Brijs et al., 2017), but higher than that reported in gill (Dawson, Millet, Selman, & Metcalfe, 2020), or muscle and liver (Salin, Auer, Anderson, Selman, & Metcalfe, 2016), though it should be noted that the protocols used in measuring Leak respiration often differ between studies. As expected, CI&CII-linked respiratory leak through the entire electron transport chain (Leak Omy) was higher than CI-linked respiratory leak (Leak CI) in our study. This confirms that it is feasible to measure both coupled and uncoupled respiration in visceral fat from Atlantic salmon.

Although we showed that about 40 mg of visceral adipose tissue, measured at 12 °C, provides a reasonable output as stated above, the demanding nature of the procedure due to logistic and physiological complexities should be noted. For example, adipose tissue quantity varies between individuals, and comparatively large tissue samples are required (previous studies have used, e.g., 8 mg from gill tissue (Dawson et al., 2020)). We also lost data points from the lowest respiration activity, i.e., three out of 16 of our measurements were unreliable for Leak CI, which could have been avoided by using a higher amount of initial material. Even more tissue would be required to repeat the measurements for each biological replicate, which was not feasible here due to taking samples for several analyses from the same fish, though previous studies in fish have found consistent mitochondrial respiration between technical replicates (Brijs et al., 2017; Dawson et al., 2020). Finally, it should be noted that since the sampled tissue can be stored only up to a few hours before the respiration measurements (unless a longer storage time is validated for this tissue (Rees, Reemeyer, & Irving, 2022)), coordination between the laboratory and the field (or rearing facilities) is demanding when research is conducted on adult-sized salmonids.

Despite the low power of the analyses due to low sample size, we detected significantly higher CI&CII-mediated respiration and CI&CII-linked respiratory leak, and marginally higher coupling capacity in immature female salmon with the *vgll3* late maturation genotype than in those with the early maturation genotype. These preliminary results may provide insight into mechanistic explanations for how variation in maturation timing is linked to *vgll3* genotype, for which we postulate below two distinct but potentially complementary hypotheses.

The first hypothesis concerns resource allocation to and from the lipid deposits in adipose tissue. Maturation in Atlantic salmon is a physiological trait with a genetic threshold mediated by condition factor possibly via lipid accumulation (Thorpe, 2007). Concordantly, a previous study (Debes et al., 2021) suggested that *vgll3*-associated early maturation in males was mediated by a higher condition factor. In line with these previous findings, and with our results, we hypothesise that the higher mitochondrial respiration in the adipose tissue of immature salmon with a late maturation genotype leads to reduced lipid storage, contributing to delayed maturation. Mitochondrial respiration typically increases during fasting – a state that is characterized by active catabolic metabolism. It is therefore tempting to speculate that the mitochondrial phenotype of the *vgll3* late maturation genotype could indicate active catabolic metabolism in adipose tissue. The high mitochondrial respiratory capacity could enhance the oxidation of energy substrates and subsequently reduce the size of adipose tissue depots in salmon carrying the *vgll3* late maturation genotype compared to the early maturation genotype. In line with this hypothesis, increased mitochondrial fatty acid oxidation in adipose tissue has been observed to lead to a lean phenotype in mice (Flachs, Rossmeisl, Kuda, & Kopecky, 2013), and reduced mitochondrial respiration is related to adipocyte hypertrophy in a cell line (Baldini et al., 2021). Further, a negative correlation between adiposity (body mass index, BMI) and respiration of isolated mitochondria from adipose tissue was observed in humans (Fischer et al., 2015). In our study, neither genotype nor mitochondrial respiration was associated with body condition of fish (analogous to BMI). A lack of *vgll3* genotype effect on body condition in salmon was also found in the same cohort as the fish we studied (Åsheim et al., 2022). However, these results do not contradict our hypothesis since body condition effects may be manifested at a different time or life-stage (Debes et al., 2021), or alternatively, was not observed due to low statistical power. For instance, salmon with the late maturation genotype could be burning their adipose tissue at a higher rate during the winter (differences in lipid utilisation have also been shown in relation to migration in juvenile salmon (Morgan, McCarthy, & Metcalfe, 2002)), which would result in faster depletion of energy reserves compared to the early maturation genotype. Subsequently, the depletion of lipid reserve could delay maturation because the amount of adipose tissue in the spring is an important determinant of salmon maturation probability the following autumn (Rowe et al., 1991).

The second hypothesis we propose is based on the finding that salmon with the *vgll3* late maturation genotype tended to have more actively working mitochondria due to a higher coupling capacity, P/E ratio. In other words, electron transfer in fish with late maturation genotype was almost maximally coupled, while coupling was only up to ∼70% in fish with the early maturation genotype. In the white adipose tissue of obese humans, coupling capacities of 77-83% (with an increasing trend during weight loss) have been observed (Hansen et al., 2015). Thus, the adipose tissue coupling capacity in salmon with the early maturation genotype was lower compared to that in obese humans, and the higher coupling capacity in salmon with late maturation genotypes is in line with active catabolic metabolism. Because coupling capacity reflects the proportion of coupled respiration from the theoretical maximum respiration, we hypothesise that it could affect the resource acquisition of salmon, especially if the effect is consistent across tissues. Specifically, the lower coupling capacity in early vs. late maturation genotype could allow the fish with early maturation genotype to increase coupled respiration more during high energy demand and changes in environmental conditions. This could happen if the remaining electron transfer capacity was used preferentially to increase ATP synthesis instead of leak respiration. The mitochondria of salmon with late maturation genotype were already near-maximally coupled despite the relatively high food availability in this study (except for a 2 d-period without feeding prior to sample collection). Thus salmon with the late maturation genotype may respond relatively poorly to increases in energy demand or stress (Sokolova, 2018), and subsequently reserve less energy available to invest into growth, maturation and ultimately survival. Such inherent high coupling capacity of salmon with the late maturation genotype may indicate that they could be more vulnerable to low food conditions such as during winters in freshwater (Mogensen & Post, 2012) or in the sea (Czorlich, Aykanat, Erkinaro, Orell, & Primmer, 2022). In line with this, a lower aerobic scope at the whole animal level – indicating lower capacity for aerobic metabolism beyond self-maintenance – in juvenile salmon with the late maturation genotype was also found by Prokkola et al., (2022). Further, the lower coupling capacity of the late maturation genotype matches the distribution of salmon with contrasting age-at-maturity in the wild, where individuals maturing later spawn typically in larger rivers that are likely more environmentally stable. To explicitly test for this hypothesis, future work should focus to detail changes in the P/E ratio by simultaneously measuring leak respiration and ATP production, which would better characterise the metabolic response, e.g., to low food availability and fasting, in salmon with different *vgll3* genotypes.

The molecular pathways that could link *vgll3* to mitochondrial respiration are not well known. In humans and mice, *vgll3* is a cofactor binding to TEA domain -containing (TEAD) transcription factors that regulates tissue differentiation pathways, such as adipogenesis and myogenesis, as well as pathways that regulate development and remodelling of tissue composition and organ and cell size, such as hippo signalling pathway (Figeac et al., 2019; Halperin et al., 2013; Hori et al., 2020). In Atlantic salmon, the widespread expression of *vgll3* is correlated with factors in the Hippo pathway (i.e. YAP, and TEAD) (Kurko et al., 2020) suggesting a functional analogy in salmon with mammals. Intriguingly, the same pathways also control mitochondrial biogenesis and function (Huang et al., 2018; Liu et al., 2020; Mammoto, Muyleart, Kadlec, Gutterman, & Mammoto, 2018), further supporting that *vgll3* genetic variation might affect mitochondrial functional variation.

We also detected sex and/or maturation effect on mtDNA amount, where (mature) males had a higher mtDNA amount (relative to gDNA) in adipose tissue than (immature) females – although there were no sex differences in mitochondrial respiration. Differences in adipose tissue processes may emerge between mature and immature individuals, because salmon consume a large part of their adipose tissue to support the high energy demand of maturation (Jonsson, Jonsson, & Hansen, 1991; Rowe et al., 1991). A previous study also suggests that sex-specific maturation schedules could be mediated by non-visceral lipid storage, e.g., in muscle (House et al., 2021). Hence, future studies that partition sex and maturation status effects across tissue types would be valuable to assess the role mitochondrial variation in relation to these two phenotypes.

## Conclusions

Adipose tissue is central to maturation as well as energy homeostasis but very little is known about how these two processes could be genetically interlinked. Our proof of principle study showed the feasibility of studying adipose tissue mitochondrial respiration in salmon and yielded insightful preliminary results from which we generated informed hypotheses for future research. To further integrate adipose tissue metabolism into a life-history evolution framework, measurements of mitochondrial respiration in salmon with different *vgll3* genotypes could be combined with analyses of lipogenesis and lipolysis (i.e., lipid synthesis and release, respectively) for example using gene expression, and lipid quantification with histochemistry. Moreover, mitochondrial respiration measurements could be combined with measurements of reactive oxygen species and the effectiveness of ATP synthesis (ATP/O) (Salin et al., 2019; Salin et al., 2018). Ultimately, to generalise the role of mitochondrial respiration and of coupling capacity in life-history evolution, studies would need to address life-stage specific genetic effects, and measurements should be extended to non-adipose tissues. Given the common physiological roles and functions of visceral white adipose tissue in salmon and humans (Salmeron, 2018), a better understanding of these functions in salmon may also facilitate its use as a new model species for obesity research. Conversely, our study provides an example of how a more advanced understanding of metabolic disorders and obesity can be harnessed to address questions relevant for ecology and evolution.

## Acknowledgements

We thank Petra Liljeström, Mikko Immonen, Paul Bangura, and numerous interns for help in fish rearing at Lammi Biological Station, the Tissue Preparation and Histochemistry Unit Meilahti, University of Helsinki, for the cryosectioning and staining, and Antti Isomäki for advice on cell size measurements. Histological Imaging was supported by HiLIFE and the Faculty of Medicine, University of Helsinki, and Biocenter Finland. The study was funded by the Academy of Finland (1328860 and 1325964 for T. A, 335443, 314383, 272376 and the Profi6 336449 funding for E.P., and 307593, 302873, 327255, and 342851 for C.R.P.), by the European Research Council under the European Articles Union’s Horizon 2020 research and innovation program (grant no. 742312), and the University of Helsinki.

## Appendices and supplemental material

Table A1. Primer details.

Supplemental material 1. Xlsx file. Table S1. The data obtained in this study. Supplemental material 2. Figures S1–S4 and Tables S3–S4.

Supplemental material 3. Txt file. A macro for adipocyte size quantification in ImageJ.

## Data availability

The final data from this study is available in Table S1. The data, R codes for analyses and figures, and the images of adipose tissue sections are available in Zenodo (https://doi.org/10.5281/zenodo.6899961).

## References

Åsheim, E. R., Debes, P. V., House, A., Niemelä, P. T., Siren, J. P., Erkinaro, J., & Primmer, C. R. (2022). Strong effects of temperature, population and age-at-maturity genotype on maturation probability for Atlantic salmon in a common garden setting. bioRxiv, 2022.2007.2022.501167. doi:10.1101/2022.07.22.501167

Baldini, F., Fabbri, R., Eberhagen, C., Voci, A., Portincasa, P., Zischka, H., & Vergani, L. (2021). Adipocyte hypertrophy parallels alterations of mitochondrial status in a cell model for adipose tissue dysfunction in obesity. Life Sciences, 265. doi:10.1016/j.lfs.2020.118812

Bankhead, P., Loughrey, M. B., Fernandez, J. A., Dombrowski, Y., McArt, D. G., Dunne, P. D., … Hamilton, P. W. (2017). QuPath: Open source software for digital pathology image analysis. Sci Rep, 7. doi:10.1038/s41598-017-17204-5

Barson, N. J., Aykanat, T., Hindar, K., Baranski, M., Bolstad, G. H., Fiske, P., … Primmer, C. R. (2015). Sex-dependent dominance at a single locus maintains variation in age at maturity in salmon. Nature, 528(7582), 405–408. doi:10.1038/nature16062

Boudina, S., & Graham, T. E. (2014). Mitochondrial function/dysfunction in white dipose tissue. Experimental Physiology, 99(9), 1168–1178. doi:10.1113/expphysiol.2014.081414

Brand, M. D. (2005). The efficiency and plasticity of mitochondrial energy transduction. Biochemical Society Transactions, 33(Pt 5), 897–904. doi:10.1042/BST0330897

Brijs, J., Sandblom, E., Sundh, H., Grans, A., Hinchcliffe, J., Ekstrom, A., … Pichaud, N. (2017). Increased mitochondrial coupling and anaerobic capacity minimizes aerobic costs of trout in the sea. Sci Rep, 7, 45778. doi:10.1038/srep45778

Canto, C., & Garcia-Roves, P. M. (2015). High-Resolution Respirometry for Mitochondrial Characterization of Ex Vivo Mouse Tissues. Current protocols in mouse biology, 5(2), 135–153. doi:10.1002/9780470942390.mo140061

Chacko, B. K., Kramer, P. A., Ravi, S., Benavides, G. A., Mitchell, T., Dranka, B. P., … Darley-Usmar, V. M. (2014). The Bioenergetic Health Index: a new concept in mitochondrial translational research. Clinical Science, 127(5-6), 367–373. doi:10.1042/cs20140101

Cousminer, D. L., Berry, D. J., Timpson, N. J., Ang, W., Thiering, E., Byrne, E. M., … Early Growth Genetics, C. (2013). Genome-wide association and longitudinal analyses reveal genetic loci linking pubertal height growth, pubertal timing and childhood adiposity. Human Molecular Genetics, 22(13), 2735–2747. doi:10.1093/hmg/ddt104

Czorlich, Y., Aykanat, T., Erkinaro, J., Orell, P., & Primmer, C. R. (2022). Rapid evolution in salmon life history induced by direct and indirect effects of fishing. Science, eabg5980. doi:10.1126/science.abg5980

Dawson, N. J., Millet, C., Selman, C., & Metcalfe, N. B. (2020). Measurement of mitochondrial respiration in permeabilized fish gills. Journal of Experimental Biology, 223(Pt 4). doi:10.1242/jeb.216762

De Pauw, A., Tejerina, S., Raes, M., Keijer, J., & Arnould, T. (2009). Mitochondrial (Dys)function in Adipocyte (De)differentiation and Systemic Metabolic Alterations. American Journal of Pathology, 175(3), 927–939. doi:10.2353/ajpath.2009.081155

Debes, P. V., Piavchenko, N., Ruokolainen, A., Ovaskainen, O., Moustakas-Verho, J. E., Parre, N., … Primmer, C. R. (2021). Polygenic and major-locus contributions to sexual maturation timing in Atlantic salmon. Molecular Ecology, 30(18), 4505–4519. doi:10.1111/mec.16062

Dennery, P. A. (2010). Oxidative stress in development: Nature or nurture? Free Radical Biology and Medicine, 49(7), 1147–1151. doi:10.1016/j.freeradbiomed.2010.07.011

Djafarzadeh, S., & Jakob, S. M. (2017). High-resolution Respirometry to Assess Mitochondrial Function in Permeabilized and Intact Cells. Jove-Journal of Visualized Experiments(120). doi:10.3791/54985

Elks, C. E., Perry, J. R., Sulem, P., Chasman, D. I., Franceschini, N., He, C., … Murray, A. (2010). Thirty new loci for age at menarche identified by a meta-analysis of genome-wide association studies. Nature Genetics, 42(12), 1077–1085. doi:10.1038/ng.714

Figeac, N., Mohamed, A. D., Sun, C., Schoenfelder, M., Matallanas, D., Garcia-Munoz, A., … Wackerhage, H. (2019). VGLL3 operates via TEAD1, TEAD3 and TEAD4 to influence myogenesis in skeletal muscle. Journal of Cell Science, 132(13). doi:10.1242/jcs.225946

Fischer, B., Schoettl, T., Schempp, C., Fromme, T., Hauner, H., Klingenspor, M., & Skurk, T. (2015). Inverse relationship between body mass index and mitochondrial oxidative phosphorylation capacity in human subcutaneous adipocytes. American Journal of Physiology-Endocrinology and Metabolism, 309(4), E380–E387. doi:10.1152/ajpendo.00524.2014

Flachs, P., Rossmeisl, M., Kuda, O., & Kopecky, J. (2013). Stimulation of mitochondrial oxidative capacity in white fat independent of UCP1: A key to lean phenotype. Biochimica Et Biophysica Acta-Molecular and Cell Biology of Lipids, 1831(5), 986–1003. doi:10.1016/j.bbalip.2013.02.003

Fleming, I. A. (1998). Pattern and variability in the breeding system of Atlantic salmon (Salmo salar), with comparisons to other salmonids. Canadian Journal of Fisheries and Aquatic Sciences, 55, 59–76. doi:DOI 10.1139/cjfas-55-S1-59

Guo, W., Jiang, L., Bhasin, S., Khan, S. M., & Swerdlow, R. H. (2009). DNA extraction procedures meaningfully influence qPCR-based mtDNA copy number determination. Mitochondrion, 9(4), 261–265. doi:10.1016/j.mito.2009.03.003

Halperin, D. S., Pan, C., Lusis, A. J., & Tontonoz, P. (2013). Vestigial-like 3 is an inhibitor of adipocyte differentiation. Journal of Lipid Research, 54(2), 473–481. doi:10.1194/jlr.M032755

Hansen, M., Lund, M. T., Gregers, E., Kraunsoe, R., Van Hall, G., Helge, J. W., & Dela, F. (2015). Adipose tissue mitochondrial respiration and lipolysis before and after a weight loss by diet and RYGB. Obesity, 23(10), 2022–2029. doi:10.1002/oby.21223

Heinonen, S., Jokinen, R., Rissanen, A., & Pietilainen, K. H. (2020). White adipose tissue mitochondrial metabolism in health and in obesity. Obes Rev, 21(2), e12958. doi:10.1111/obr.12958

Hood, W. R., Austad, S. N., Bize, P., Jimenez, A. G., Montooth, K. L., Schulte, P. M., … Salin, K. (2018). The Mitochondrial Contribution to Animal Performance, Adaptation, and Life-History Variation INTRODUCTION. Integrative and Comparative Biology, 58(3), 480–485. doi:10.1093/icb/icy089

Hori, N., Okada, K., Takakura, Y., Takano, H., Yamaguchi, N., & Yamaguchi, N. (2020). Vestigial-like family member 3 (VGLL3), a cofactor for TEAD transcription factors, promotes cancer cell proliferation by activating the Hippo pathway. Journal of Biological Chemistry, 295(26), 8798–8807. doi:10.1074/jbc.RA120.012781

House, A. H., Debes, P. V., Kurko, J., Erkinaro, J., Kakela, R., & Primmer, C. R. (2021). Sex-specific lipid profiles in the muscle of Atlantic salmon juveniles. Comparative Biochemistry and Physiology D-Genomics & Proteomics, 38. doi:10.1016/j.cbd.2021.100810

Huang, S., Wang, X., Wu, X., Yu, J., Li, J., Huang, X., … Ge, H. (2018). Yap regulates mitochondrial structural remodeling during myoblast differentiation. American Journal of Physiology: Cell Physiology, 315(4), C474–C484. doi:10.1152/ajpcell.00112.2018

Jokinen, R., Pirnes-Karhu, S., Pietilainen, K. H., & Pirinen, E. (2017). Adipose tissue NAD(+)-homeostasis, sirtuins and poly(ADP-ribose) polymerases - important players in mitochondrial metabolism and metabolic health. Redox Biology, 12, 246–263. doi:10.1016/j.redox.2017.02.011

Jonsson, N., Jonsson, B., & Hansen, L. P. (1991). Energetic cost of spawning in male and female Atlantic salmon (Salmo salar L). Journal of Fish Biology, 39(5), 739–744. doi:10.1111/j.1095-8649.1991.tb04403.x

Kadenbach, B. (2003). Intrinsic and extrinsic uncoupling of oxidative phosphorylation. Biochimica Et Biophysica Acta-Bioenergetics, 1604(2), 77–94. doi:10.1016/s0005-2728(03)00027-6

Koch, R. E., Buchanan, K. L., Casagrande, S., Crino, O., Dowling, D. K., Hill, G. E., … Stier, A. (2021). Integrating Mitochondrial Aerobic Metabolism into Ecology and Evolution. Trends in Ecology & Evolution, 36(4), 321–332. doi:10.1016/j.tree.2020.12.006

Kurko, J., Debes, P. V., House, A. H., Aykanat, T., Erkinaro, J., & Primmer, C. R. (2020). Transcription Profiles of Age-at-Maturity-Associated Genes Suggest Cell Fate Commitment Regulation as a Key Factor in the Atlantic Salmon Maturation Process. G3-Genes Genomes Genetics, 10(1), 235–246. doi:10.1534/g3.119.400882

Lailvaux, S. P., & Husak, J. F. (2014). The life-history of whole-organism performance. Quarterly Review of Biology, 89(4), 285–318. doi:10.1086/678567

Liu, R., Jagannathan, R., Sun, L., Li, F., Yang, P., Lee, J., … Moulik, M. (2020). Tead1 is essential for mitochondrial function in cardiomyocytes. American Journal of Physiology: Heart and Circulatory Physiology, 319(1), H89–H99. doi:10.1152/ajpheart.00732.2019

Mammoto, A., Muyleart, M., Kadlec, A., Gutterman, D., & Mammoto, T. (2018). YAP1-TEAD1 signaling controls angiogenesis and mitochondrial biogenesis through PGC1alpha. Microvasc Res, 119, 73–83. doi:10.1016/j.mvr.2018.04.003

Martin, A. S. G., & Obin, M. S. (2006). Obesity and the role of adipose tissue in inflammation and metabolism. American Journal of Clinical Nutrition, 83(2), 461S–465S. doi:10.1093/ajcn/83.2.461s

Mobley, K. B., Aykanat, T., Czorlich, Y., House, A., Kurko, J., Miettinen, A., … Primmer, C. R. (2021). Maturation in Atlantic salmon (Salmo salar, Salmonidae): a synthesis of ecological, genetic, and molecular processes. Reviews in Fish Biology and Fisheries, 31(3), 523–571. doi:10.1007/s11160-021-09656-w

Mogensen, S., & Post, J. R. (2012). Energy allocation strategy modifies growth-survival trade-offs in juvenile fish across ecological and environmental gradients. Oecologia, 168(4), 923–933. doi:10.1007/s00442-011-2164-0

Mohamed-Ali, V., Pinkney, J. H., & Coppack, S. W. (1998). Adipose tissue as an endocrine and paracrine organ. International Journal of Obesity, 22(12), 1145–1158. doi:10.1038/sj.ijo.0800770

Morgan, I. J., McCarthy, I. D., & Metcalfe, N. B. (2002). The influence of life-history strategy on lipid metabolism in overwintering juvenile Atlantic salmon. Journal of Fish Biology, 60(3), 674–686. doi:10.1006/jfbi.2002.1886

Nanton, D. A., Vegusdal, A., Rørå, A. M. B., Ruyter, B., Baeverfjord, G., & Torstensen, B. E. (2007). Muscle lipid storage pattern, composition, and adipocyte distribution in different parts of Atlantic salmon (Salmo salar) fed fish oil and vegetable oil. Aquaculture, 265(1-4), 230–243. doi:10.1016/j.aquaculture.2006.03.053

Norgan, N. G. (1997). The beneficial effects of body fat and adipose tissue in humans. Int J Obes Relat Metab Disord, 21(9), 738–746. doi:10.1038/sj.ijo.0800473

Ottaviani, E., Malagoli, D., & Franceschi, C. (2011). The evolution of the adipose tissue: A neglected enigma. General and Comparative Endocrinology, 174(1), 1–4. doi:10.1016/j.ygcen.2011.06.018

Pfaffl, M. W. (2001). A new mathematical model for relative quantification in real-time RT-PCR. Nucleic Acids Research, 29, -.

Prokkola, J. M., Asheim, E. R., Morozov, S., Bangura, P., Erkinaro, J., Ruokolainen, A., … Aykanat, T. (2022). Genetic coupling of life-history and aerobic performance in Atlantic salmon. Proc Biol Sci, 289(1967), 20212500. doi:10.1098/rspb.2021.2500

R Core Team. (2019). R: A language and environment for statistical computing. In: R Foundation for Statistical Computing, Vienna, Austria.

Rees, B. B., Reemeyer, J. E., & Irving, B. A. (2022). Interindividual variation in maximum aerobic metabolism varies with gill morphology and myocardial bioenergetics in Gulf killifish. Journal of Experimental Biology, 225(12). doi:10.1242/jeb.243680

Robin, E. D., & Wong, R. (1988). Mitochondrial-DNA molecules and virtual number of mitochondria per cell in mammalian cells. Journal of Cellular Physiology, 136(3), 507–513. doi:10.1002/jcp.1041360316

Rowe, D. K., Thorpe, J. E., & Shanks, A. M. (1991). Role of fat stores in the maturation of male Atlantic salmon (Salmo salar) parr. Canadian Journal of Fisheries and Aquatic Sciences, 48(3), 405–413. doi:10.1139/f91-052

Ruijter, J. M., Ramakers, C., Hoogaars, W. M. H., Karlen, Y., Bakker, O., van den Hoff, M. J. B., & Moorman, A. F. M. (2009). Amplification efficiency: linking baseline and bias in the analysis of quantitative PCR data. Nucleic Acids Research, 37, -.

Salin, K., Auer, S. K., Anderson, G. J., Selman, C., & Metcalfe, N. B. (2016). Inadequate food intake at high temperatures is related to depressed mitochondrial respiratory capacity. Journal of Experimental Biology, 219(9), 1356–1362. doi:10.1242/jeb.133025

Salin, K., Auer, S. K., Rey, B., Selman, C., & Metcalfe, N. B. (2015). Variation in the link between oxygen consumption and ATP production, and its relevance for animal performance. Proc Biol Sci, 282(1812), 20151028. doi:10.1098/rspb.2015.1028

Salin, K., Auer, S. K., Rudolf, A. M., Anderson, G. J., Selman, C., & Metcalfe, N. B. (2016). Variation in Metabolic Rate among Individuals Is Related to Tissue-Specific Differences in Mitochondrial Leak Respiration. Physiological and Biochemical Zoology, 89(6), 511–523. doi:10.1086/688769

Salin, K., Villasevil, E. M., Anderson, G. J., Lamarre, S. G., Melanson, C. A., McCarthy, I., … Metcalfe, N. B. (2019). Differences in mitochondrial efficiency explain individual variation in growth performance. Proceedings of the Royal Society Biological Sciences Series B, 286(1909), 1–8.

Salin, K., Villasevil, E. M., Anderson, G. J., Selman, C., Chinopoulos, C., & Metcalfe, N. B. (2018). The RCR and ATP/O Indices Can Give Contradictory Messages about Mitochondrial Efficiency. Integrative and Comparative Biology, 58(3), 486–494. doi:10.1093/icb/icy085

Salmeron, C. (2018). Adipogenesis in fish. Journal of Experimental Biology, 221(Pt Suppl 1). doi:10.1242/jeb.161588

Sinclair-Waters, M., Odegard, J., Korsvoll, S. A., Moen, T., Lien, S., Primmer, C. R., & Barson, N. J. (2020). Beyond large-effect loci: large-scale GWAS reveals a mixed large-effect and polygenic architecture for age at maturity of Atlantic salmon. Genetics Selection Evolution, 52(1), 9. doi:10.1186/s12711-020-0529-8

Sokolova, I. (2018). Mitochondrial Adaptations to Variable Environments and Their Role in Animals’ Stress Tolerance. Integrative and Comparative Biology, 58(3), 519–531. doi:10.1093/icb/icy017

Thorpe, J. E. (2007). Maturation responses of salmonids to changing developmental opportunities. Marine Ecology Progress Series, 335, 285–288. doi:10.3354/meps335285

Wickham, H. (2009). ggplot2: Elegant Graphics for Data Analysis. Ggplot2: Elegant Graphics for Data Analysis, 1–212. doi:10.1007/978-0-387-98141-3

Ye, J., Coulouris, G., Zaretskaya, I., Cutcutache, I., Rozen, S., & Madden, T. L. (2012). Primer-BLAST: A tool to design target-specific primers for polymerase chain reaction. BMC Bioinformatics, 13. doi:10.1186/1471-2105-13-134

