## Supplementary material for "Adipose tissue mitochondrial respiration in Atlantic salmon: implications for sex-dependent life-history variation": Table A1

**Table A1:** Specification of Primers, qPCR efficiencies, and Ct values for four markers used in the study.

| Gene name | Primer name | Gene symbol | Forward 5->3 | Reverse 5->3 | Product (bp) | Chr. | Pos (start) | qPCR eff (%) | Ct (mean) | Ct (sd) |
|---|---|---|---|---|---|---|---|---|---|---|
| Translation elongation factor 1 alpha | ss_ef1a_2[1,2] | *EF1a* | GGGCAGTAGTGTGGATTCCTT | CCTAGGGTGTTGATTGTGGCAA | 109 | 1 | 58223687 | 90.3 | 28.77 | 1.30 |
| Amyloid beta A4 protein-like | ss_app_1[1,3] | *App* | CATGTGGGACCGTATGGCTT | GGCATGGAGCTCAGCCTTAT | 108 | 16 | 61003412 | 89.2 | 29.11 | 1.37 |
| 16S ribosomal protein | ss_mtDNA_16S_1[4] | *16s* | ATGGCATCACGAGGGCTTAG | CTTGACGTGATCTGCCTGGT | 131 | mtDNA | 3142 | 90.6 | 22.24 | 1.56 |
| Cytochrome b | ss_mtDNA_cytb_3[4] | *Cytb* | ACGCAATCCTACGCTCCATT | AATTGGGTGAGTGGGCGAAA | 136 | mtDNA | 16215 | 91.4 | 20.78 | 1.45 |

1 GCF_000233375.1_ICSASG_v2, *Salmo salar* genome (Lien et al., 2016)

2 LOC106574335, Translation elongation factor 1 alpha

3 LOC106607006, amyloid beta A4 protein-like

4 *Salmo salar* mitochondrion, complete genome (NC_001960.1) (Feng et al., 2017)

### References


Feng, J., Wu, Z., Xie, X., Dai, Z., & Liu, S. (2017). A real-time polymerase chain reaction method for the identification of four commercially important salmon and trout species. *Mitochondrial DNA Part A*, *28*, 104-111. https://doi.org/10.3109/19401736.2015.1111346

Lien, S., Koop, B. F., Sandve, S. R., Miller, J. R., Kent, M. P., Nome, T., . . . Davidson, W. S. (2016). The Atlantic salmon genome provides insights into rediploidization. *Nature*, *533*, 200-205. https://doi.org/10.1038/nature17164
