## Supplemental material 2 for "Adipose tissue mitochondrial respiration in Atlantic salmon: implications for sex-dependent life-history variation"

### Supplemental Figures

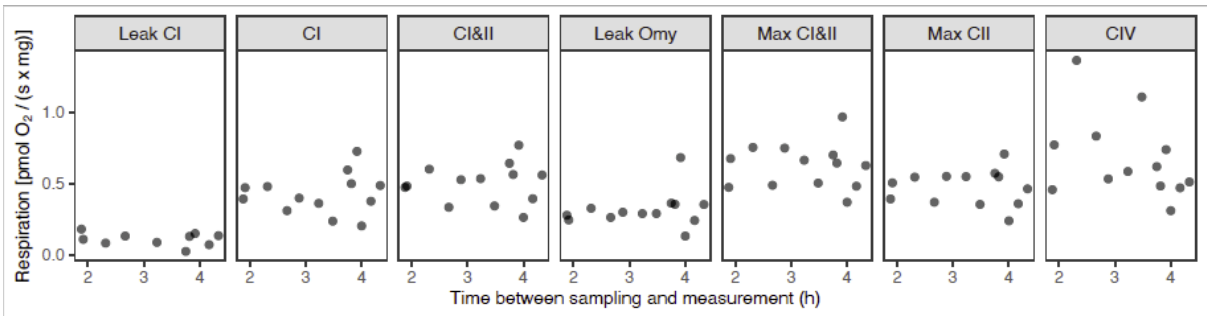


Fig. S1. Scatter plots of respiration fluxes and time lag between sampling and respiration measurement. All Spearman's rank correlation *rho* were non-significant ($p > 0.17$).

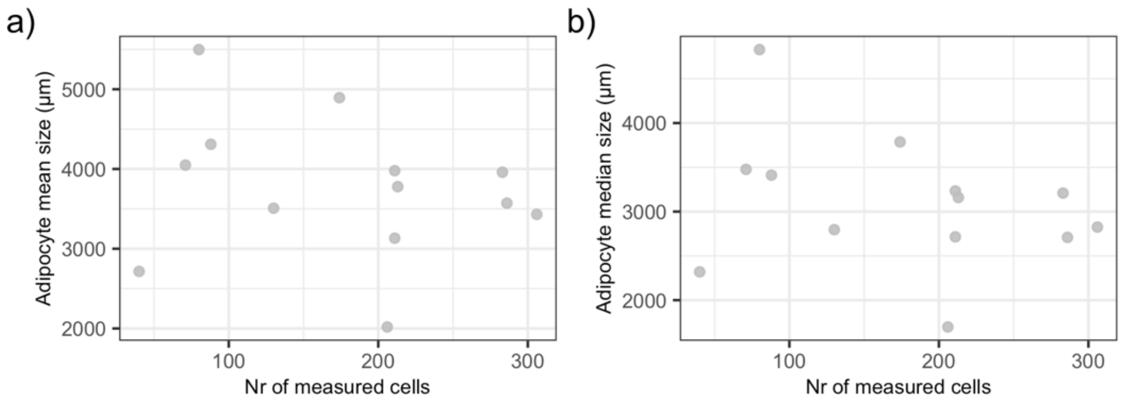


Fig. S2. The (a) mean and (b) median of adipocyte area vs. the number of cells measured from each individual (shown by grey points).

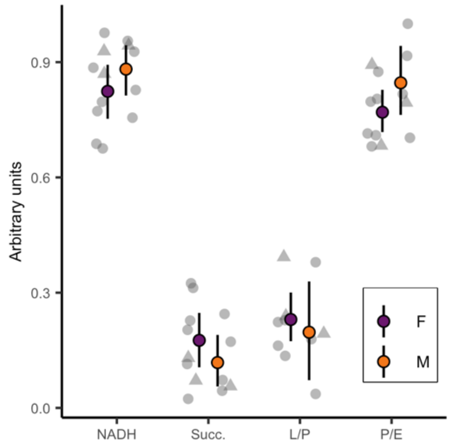


Fig. S3. Sex-specific respiration coefficients. No significant differences between sexes (Wilcoxon rank sum tests: NADH W = 14, p = 0.421; Succ. W = 26, p = 0.421; L/P W = 15, p = 0.594; P/E W = 12, p = 0.272). Coloured points show means ± 95% bootstrapped confidence intervals. Grey symbols show individual data points: circles = Control, triangles = Low fat.

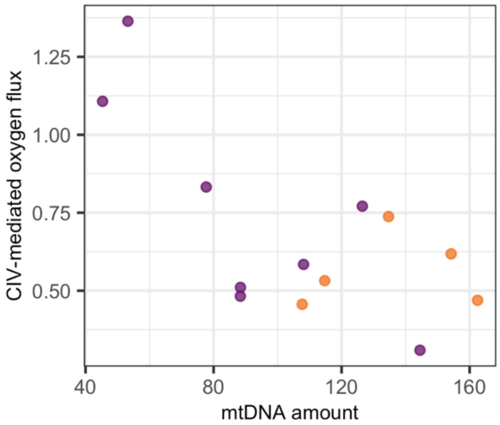


Fig. S4. Scatter plot of CIV-mediated oxygen flux (pmol $O_2$ / s*mg) and relative mtDNA amount in adipose tissue. Colours indicating sex (purple = female, orange = male).

**Supplemental tables**

Tables S1-S2 in separate files.

Table S**3**. Results of Wilcoxon rank sum tests for feed treatment effects on mitochondrial respiration. Groups were combined as no differences were detected.

| | W | p |
|---|---|---|
| Leak CI | 3 | 0.111 |
| CI | 10 | 0.447 |
| CI & CII | 12 | 0.673 |
| Leak Omy | 9 | 0.353 |
| Max CI & CII | 13 | 0.8 |
| Max CII | 13 | 0.8 |
| CIV | 11 | 0.554 |

Table S4. Pairwise Spearman's rank correlations between adipose tissue and fish phenotypes with mitochondrial respiration parameters in Atlantic salmon. See CIV and mtDNA in Fig. S4.

| **Variables** | | **Spearman *rho*** | **p** |
|---|---|---|---|
| **Adipocyte size** | CI | -0.318 | 0.341 |
| | CI&CII | -0.245 | 0.468 |
| | CIV | 0.109 | 0.755 |
| | Leak Omy | -0.255 | 0.451 |
| | Leak CI | -0.429 | 0.299 |
| | Max CI&CII | -0.118 | 0.734 |
| | Max CII | -0.245 | 0.468 |
| **mtDNA** | CI | 0.154 | 0.617 |
| | CI&CII | 0.066 | 0.835 |
| | CIV | -0.495 | 0.089 |
| | Leak Omy | -0.176 | 0.566 |
| | Leak CI | -0.309 | 0.387 |
| | Max CI&CII | -0.005 | 0.993 |
| | Max CII | 0.187 | 0.541 |
| **Fish body mass** | CI | 0.022 | 0.949 |
| | CI&CII | -0.099 | 0.751 |
| | CIV | 0.159 | 0.604 |
| | Leak Omy | -0.121 | 0.696 |
| | Leak CI | -0.164 | 0.657 |
| | Max CI&CII | -0.016 | 0.964 |
| | Max CII | -0.176 | 0.566 |
| **Fish condition factor** | CI | -0.220 | 0.470 |
| | CI&CII | -0.319 | 0.289 |
| | CIV | -0.242 | 0.426 |
| | Leak Omy | -0.291 | 0.334 |
| | Leak CI | -0.491 | 0.154 |
| | Max CI&CII | -0.335 | 0.263 |
| | Max CII | -0.352 | 0.239 |
